# Global analysis of the specificities and targets of endoribonucleases from *E. coli* toxin-antitoxin systems

**DOI:** 10.1101/2021.07.08.451724

**Authors:** Peter H. Culviner, Isabel Nocedal, Sarah M. Fortune, Michael T. Laub

**Affiliations:** Department of Immunology and Infectious Diseases, Harvard T.H. Chan School of Public Health Boston, Massachusetts USA; Department of Biology Massachusetts Institute of Technology Cambridge, MA 02139 USA; Howard Hughes Medical Institute Massachusetts Institute of Technology Cambridge, MA 02139 USA

## Abstract

Toxin-antitoxin systems are widely distributed genetic modules typically featuring toxins that can inhibit bacterial growth and antitoxins that can reverse inhibition. Although *Escherichia coli* encodes 11 toxins with known or putative endoribonuclease activity, the target of most of these toxins remain poorly characterized. Using a new RNA-seq pipeline that enables the mapping and quantification of RNA cleavage with single-nucleotide resolution, we characterize the targets and specificities of 9 endoribonuclease toxins from *E. coli*. We find that these toxins use low-information cleavage motifs to cut a significant proportion of mRNAs in *E. coli*, but not tRNAs or the rRNAs from mature ribosomes. However, all the toxins, including those that are ribosome-dependent and cleave only translated RNA, inhibit ribosome biogenesis. This inhibition likely results from the cleavage of ribosomal protein transcripts, which disrupts the stoichiometry and biogenesis of new ribosomes and causes the accumulation of aberrant ribosome precursors. Collectively, our results provide a comprehensive, global analysis of endoribonuclease-based toxin-antitoxin systems in *E. coli* and support the conclusion that, despite their diversity, each disrupts translation and ribosome biogenesis.

**Importance:** Toxin-antitoxin systems are widespread genetic modules found in almost all bacteria that can regulate their growth and may play prominent roles in phage defense. *Escherichia coli* encodes 11 TA systems in which the toxin is a known or predicted endoribonuclease. The targets and cleavage specificities of these endoribonucleases have remained largely uncharacerized, precluding an understanding of how each impacts cell growth and an assessment of whether they have distinct or overlapping targets. Using a new and broadly applicable RNA-seq pipeline, we present a global analysis of 9 endoribonulease toxins from *E. coli*. We find that each uses a relatively low-information cleavage motif to cut a large proportion of mRNAs in *E. coli*, but not tRNAs or mature rRNAs. Notably, although the precise set of targets varies, each toxin efficiently disrupts ribosome biogenesis, primarily by cleaving the mRNAs of ribosomal proteins. In sum, the analyses presented provide new, comprehensive insights into the cleavage specificities and targets of almost all endoribonuclease toxins in *E. coli*. Despite different specificities, our work reveals a striking commonality in function as each toxin disrupts ribosome biogenesis and translation.

## Introduction

Toxin-antitoxin (TA) systems are genetic modules distributed across bacteria and archaea that can regulate or inhibit the growth of their host cell^1–3^. Although first characterized as plasmid maintenance systems, most TA systems are encoded on bacterial chromosomes. There are multiple types of TA systems, classified based on the nature of the antitoxin. One of the most common types is type II TA systems. These encode a toxin that inhibits growth and a co-expressed antitoxin protein that binds and inhibits the toxin. Under conditions that remain poorly defined, antitoxins are thought to be degraded or liberated from their cognate toxins, thereby freeing active toxin to inhibit growth^1^. The toxins of TA systems comprise several different protein families, with a diversity of mechanisms for inhibiting cell growth^2, 4^.

TA systems have been suggested to promote adaptation to various stresses^3^. The ectopic expression of many toxins causes cells to enter a growth-arrested state in which they are stress- and antibiotic-tolerant^5–7^. Additionally, the transcription of many TA systems increases in response to a range of stresses^8^. However, there are few cases of strong, reproducible deletion phenotypes for TA systems and recent work has demonstrated that toxins may not be activated even if their transcription is induced by a stress condition^9, 10^. TA systems were also suggested to contribute to spontaneous persister cell formation, but at least for *E. coli*, this has been refuted^9, 11^. Some TA systems are activated during phage infection, with the toxin leading to abortive infection, either by inhibiting host cell processes or disrupting phage replication or maturation^12–14^.

Many bacteria encode endoribonuclease toxins that have been shown to cleave a variety of RNAs, including mRNAs, tRNAs, and rRNAs^15–20^. These endoribonucleases can be ribosome-independent or ribosome-dependent, with the latter class requiring an interaction with the ribosome to reposition catalytic amino acids that then drive cleavage of translated mRNAs, often at a particular position within codons^16, 21, 22^. *E. coli* MG1655 (hereafter *E. coli* for simplicity) encodes 11 type II TA systems where the toxin is known or predicted to have endoribonuclease activity, but the precise targets and specificities of most of these toxins remains unclear (Table 1). Six of these toxins have been suggested to be ribosome-dependent and thus require translation of their target RNAs.

**Table 1.** Summary of prior literature on toxin specificity.

| **Toxin** | **Family or Superfamily** | **Ribosome Dependence** | **Core Cleavage Specificity** |
| --- | --- | --- | --- |
| MazF | MazF<sup>4</sup> | No<sup>19,39</sup> | ^ACA[AUC]<sup>19,39</sup> |
| ChpB | MazF<sup>4,40</sup> | No<sup>40</sup> | ^A^C[UGC]<sup>40</sup> |
| RnlA | RnlA, sometimes fused to RNase HI domains<sup>41</sup> | Unknown<sup>41</sup> | Poorly defined<sup>41</sup> |
| HicA | HicA<sup>27</sup> | No<sup>27</sup> | Poorly defined<sup>27</sup> |
| MqsR | RelE (structural homology)<sup>31,32</sup>, GinB<sup>4</sup> | No<sup>17,20,42</sup>, possible ribosome interaction<sup>32</sup> | G^CU<sup>17,20,42</sup> |
| RelE | RelE<sup>4</sup> | Yes<sup>16</sup> | C^G<sup>16</sup>, coding frame preference 12^3<sup>16</sup> |
| YhaV | RelE<sup>43</sup> | Yes<sup>44</sup>, No<sup>43</sup> | Poorly defined<sup>44</sup> |
| HigB | RelE<sup>4</sup> | Yes<sup>31</sup> | Poorly defined<sup>31</sup> |
| YoeB | RelE<sup>4</sup> | Yes<sup>45</sup> | ^[AG]<sup>46</sup> |
| YafO | YafO<sup>4</sup>, RelE (weak sequence and structural homology)<sup>31</sup> | Yes<sup>31</sup> | Poorly defined<sup>31</sup> |
| YafQ | RelE<sup>4</sup> | Yes<sup>47,48</sup> | AA^A<sup>48</sup>, ^A<sup>47</sup>; coding frame preference 12^3<sup>47,48</sup> |

Previously, we developed a quantitative RNA-Seq-based approach to systematically map the RNA targets of *E. coli* MazF, a ribosome-independent toxin. Inducing MazF leads to rapid, widespread cleavage of mRNAs, producing a global disruption in translation, consistent with earlier studies concluding that MazF is an mRNA interferase^19, 23^. Additionally, we found that MazF drives the accumulation of rRNA precursors, likely by directly cleaving nascent rRNA and by cleaving ribosomal protein transcripts to prevent their translation, leading to defects in rRNA processing and ribosome assembly. These results suggested that MazF can inhibit cell growth by targeting many mRNAs and inhibiting the proper synthesis of ribosomes. This mechanism of growth inhibition is enabled in part by MazF’s highly abundant cleavage site, the trinucleotide ACA, with some additional specificity in the two nucleotides on either side of the ACA. Like MazF, the toxin MqsR also directly cleaves rRNA precursors^20^. Whether other toxins are also able to disrupt ribosome biogenesis has, to our knowledge, not been systematically investigated.

To characterize the endoribonuclease toxins in *E. coli* and compare their specificities and targets, we developed a new RNA-seq pipeline that enabled the identification and quantification of RNA cleavage events with single-nucleotide resolution. We found that each toxin recognizes a short, low complexity motif and, consequently, cleaved much of the transcriptome after induction. Ribosome-independent toxins show no bias toward cleaving a particular codon position and generally cleave throughout the coding region of target transcripts. In contrast, ribosome-dependent toxins showed clear bias to cleaving near the 5’-ends of translated mRNAs and at specific locations within subcodons. Our results support the ribosome-dependency suggested previously for most toxins but suggest a reconsideration of the ribosome-dependency of HicA, YafO, and MqsR. For all the toxins, we found no evidence for the direct cleavage of mature rRNAs or tRNAs. However, each inhibited ribosome biogenesis leading to the accumulation of abnormal rRNA precursors. Importantly, this inhibition of ribosome biogenesis occurred even for ribosome-dependent toxins that cannot cut rRNA precursors directly, strongly favoring the model that toxins indirectly disrupt ribosome biogenesis through decreased translation of ribosomal protein transcripts. These results suggest that through short, low information content motifs each endoribonuclease toxin in *E. coli* can efficiently disrupt the proper synthesis of ribosomes and likely other large multiprotein complexes to inhibit growth.

## Results

### Most toxins inhibit cell growth and can be antagonized by a cognate antitoxin

To compare the effects of *E. coli*’s toxins on the transcriptome, we first cloned each toxin and antitoxin into a common expression system. Using separate but compatible low-copy plasmids in a wild-type *E. coli* MG1655 background, we expressed each toxin from an arabinose-inducible promoter and each cognate antitoxin from a tetracycline-inducible promoter. Strains harboring each toxin-antitoxin pair were grown to mid-exponential phase (OD_600_ ∼ 0.25-0.3) in M9 glycerol, followed by back dilution to OD_600_∼0.1 with the addition of either arabinose or arabinose and anhydrotetracycline to induce the toxin alone or the toxin and antitoxin, respectively (Figure 1A). Growth from 30 minutes to 2 hours after induction was used to measure the growth rate. For 8 of 11 TA systems, we observed a greater than 2-fold increase in doubling time when inducing the toxin alone compared to both the toxin and antitoxin (Figure 1B). For HicA, there was a small, but not significant change in doubling time when expressing the toxin alone, possibly because leaky expression of the antitoxin was sufficient to largely neutralize the toxin. For YafQ and RnlA, induction of the toxin alone did not substantially increase doubling time (Figure S1A). Because these experiments were conducted in a wild-type background, the endogenous copies of antitoxin (*dinJ* and *rnlB*, respectively) may have been sufficient to inhibit toxicity^13^. In subsequent analyses, we focused on the 9 systems (Figure 1B) where the toxin alone affected growth rate and the antitoxin could rescue the growth defect.

**Figure 1:**
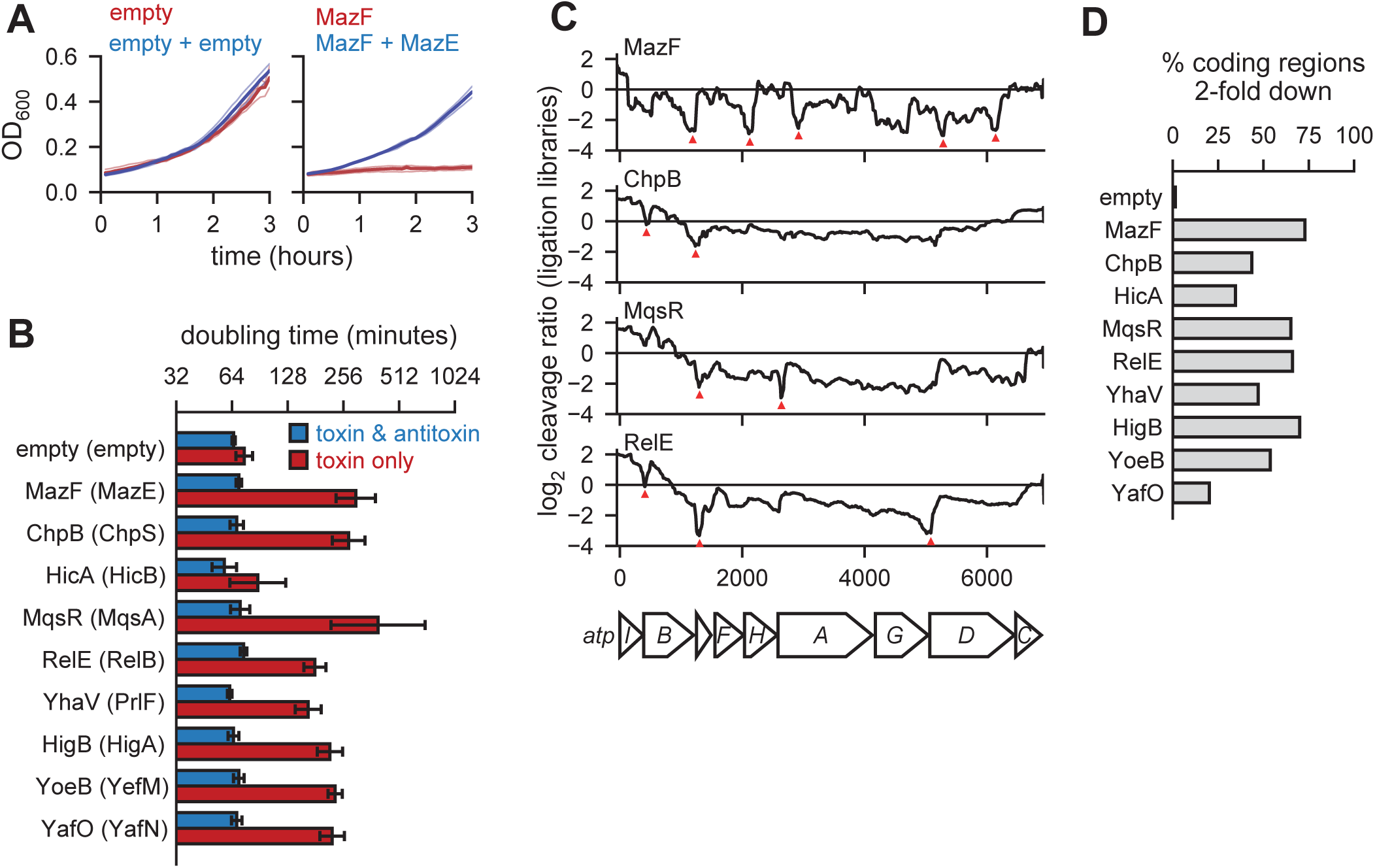
Endoribonuclease toxins inhibit cell growth and broadly cleave the transcriptome. **(A)** Sample growth data for cells harboring two empty vectors (left) or toxin (MazF) and antitoxin (MazE) containing vectors. Toxin was induced (red) or toxin and antitoxin were co-induced (blue) at t=0. Three replicates (faded lines) were used to calculate the mean (solid lines). **(B)** Doubling times calculated from growth curves like those in (A) using data between 30 minutes and 2 hours for cells expressing a given toxin (red) or toxin and antitoxin (blue). Error bars show the standard deviation of 3 replicates. **(C)** Cleavage profiles of the highly expressed *atpI-C* region are plotted for four toxins. Selected narrow valleys are highlighted with red triangles. All toxins were expressed for 10 minutes with the exception of MazF, which was expressed for 5 minutes. Data shown is from a single ligation RNA-seq library for each toxin compared to a single empty vector library. **(D)** The percentage of highly-expressed (all positions ≥64 reads in the vector control) coding regions in *E. coli* with at least one position having a cleavage ratio below -1, or a relative 2-fold downregulation of RNA abundance at the site, after expressing each toxin indicated. Data are from ligation libraries as in Figure 1C.

### Toxin expression results in widespread degradation of *E. coli* transcripts

To determine how toxins affect *E. coli* mRNAs, we conducted strand-specific paired-end RNA-seq after expressing each toxin for 10 minutes and mapped the full nucleotide coverage of the resulting reads. As in our prior study of MazF^19^, we calculated a cleavage ratio (log_2_ + toxin reads : empty-vector reads) at each nucleotide across transcripts. A negative cleavage ratio indicates that a region is either cleaved or destabilized following induction of a given toxin. We examined cleavage ratios across transcripts, hereafter called cleavage profiles (Figure 1C). Endoribonuclease activity generates valleys within cleavage profiles, whereas changes in general expression or stability causes a transcript’s entire cleavage profile to increase or decrease. Visual inspection of the cleavage profiles for the toxins revealed several patterns: (i) the 9 toxins each generated discrete valleys within some transcripts, supporting the notion that each has endoribonucleolytic activity; (ii) cleavage profiles, and the position of the valleys, differed depending on which toxin was induced, implying that the toxins have different cleavage specificities; and (iii) cleavage ratio values often decreased well below 0, indicating that toxin expression led to substantial amounts of cleavage. Apart from YafO and HicA, toxin expression led to a minimum cleavage ratio < −1 (*i.e.* > 2-fold down) in more than 40% of all expressed coding regions (Figure 1D). Together, these observations support the conclusion that toxin expression results in a major remodeling of the *E. coli* transcriptome through direct cleavage of mRNA at a wide variety of sites.

### Mapping and quantification of endoribonuclease-dependent cleavage

To better understand the effect of each toxin on the transcriptome, we initially attempted to extract cleavage specificity from cleavage valleys using an approach analogous to our previous work on MazF^19^. However, this approach was complicated by the broad and shallow valleys generated by many of the other toxins compared to MazF (Figure 1C); as the size of a valleys increases it becomes difficult to identify common sequences between valleys and thus determine a sequence specificity. Other methods, based on enrichment and sequencing of RNAs containing the 5’-OH left by many endoribonuclease toxins, can pinpoint cleavage sites but are not quantitative^18^. We therefore developed a new, dual library construction protocol to achieve both single-nucleotide resolution of cleavage sites and quantification of the extent of cleavage, while also increasing the throughput of library construction compared prior approaches (Figure 2A). Briefly, to identify 5’-ends produced from cleavage events, we used random primers containing a sample-specific barcode and PCR handle at their 5’-ends to initiate cDNA synthesis using an MMLV-type reverse transcriptase (RT). Upon transcribing to the end of an RNA molecule, this RT enzyme will add non-templated cytosines that can then hybridize to the guanosines at the 3’-end of a template switching oligo, which encodes a second PCR handle (Figure 2B). The resulting cDNAs can then be amplified by PCR and sequenced. The overall approach is similar to existing low-input and single-cell sequencing techniques^24, 25^. By sequencing with a custom primer matching the template switching oligo, we can pinpoint the 5’-end of the RNAs generated by each endoribonuclease toxin (Figure 2C).

**Figure 2:**
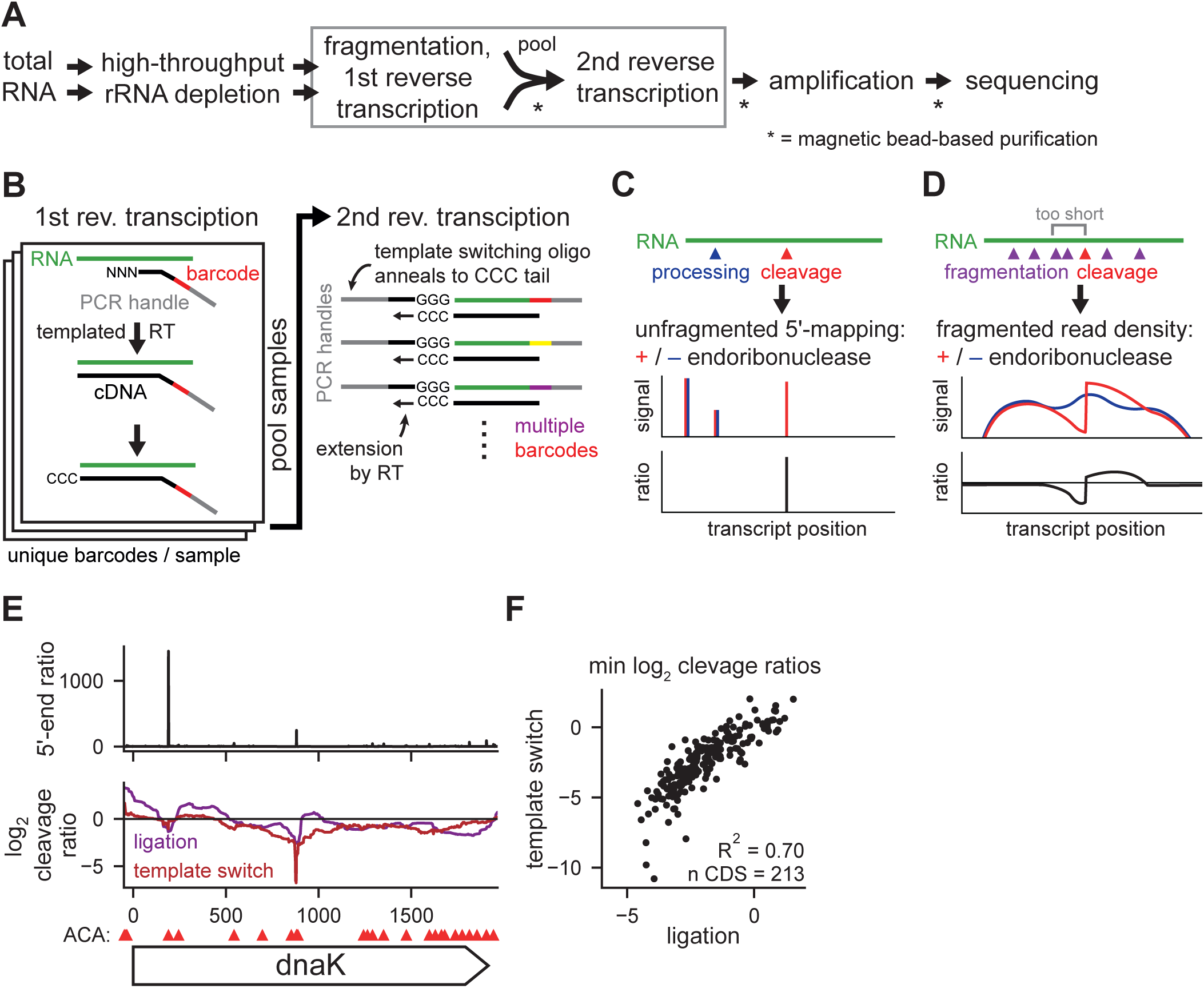
Using template switch libraries to characterize endoribonuclease cleavage. **(A)** Workflow for template switching libraries. Steps with multiple arrows in between are un-pooled and single arrow steps are pooled. Steps requiring a purification are starred. Steps shown in (B) are boxed. **(B)** Cartoon illustrating the molecular details of the 1^st^ and 2^nd^ reverse transcription steps in (A). **(C)** Cartoon illustrating how 5’-ends are mapped at single-nucleotide resolution using template switch libraries. **(D)** Cartoon illustrating how the extent of cleavage is quantified using fragmented template switch libraries. **(E)** Mapping of MazF cleavage across the *dnaK* gene after 5 minutes of toxin expression. (Top) 5’-end ratios from template switch libraries (geometric mean of 2 replicates). (Middle) Cleavage profiles from fragmented template switching libraries (red, n=2) and ligation libraries (purple, n=2). (Bottom) ACA sites within *dnaK* are shown as red triangles above the gene. **(F)** Scatter plot comparing coding region minimum cleavage ratios from template switch and ligation libraries within highly expressed coding regions (≥ 256 reads at all positions in both library types).

To quantify cleavage we measured the loss of reads near cleavage sites as the site of cleavage cannot be crossed by RT during cDNA synthesis. However, because our technique relies on a free 5’-end for an RNA to be detected, 3’-regions of long, intact transcripts would be under-represented in our sequencing. Thus, to generate 5’-ends throughout transcripts we fragmented the input RNA prior to library construction. The resulting cleavage profile shows a loss of read density immediately upstream of a cleavage site, but not necessarily downstream as cleavage by the toxin may also create a stable, sequenceable 5’-end (Figure 2D). In sum, our new method involves two libraries from the same initial rRNA-depleted sample, one that allows mapping of cleavage sites with single-nucleotide resolution and one that provides quantification of cleavage at each site. For simplicity, we will refer to these libraries as unfragmented and fragmented template switch libraries, respectively.

To validate this new method, we characterized the cleavage sites of MazF, whose cleavage specificity and targets we had already mapped in a prior study using the adapter ligation-based RNA-seq protocol we used for our initial toxin expression data (Figure 1C)^19^; for simplicity we refer to this as a ligation library. For both the template switch and ligation libraries, we expressed the MazF from a low-copy plasmid for 5 minutes and compared to an empty vector control. For the representative transcript *dnaK*, MazF cleavage profiles derived from the ligation library identified two cleavage valleys at positions ∼200 and ∼900 nt (Figure 2E, bottom). The unfragmented template switch library produced two major peaks within *dnaK* at the same two sites (Figure 2E, top). The corresponding fragmented template switch library produced a cleavage profile with valleys at these two sites (Figure 2E, bottom). The first site had a minimum cleavage ratio of -1 while the second site had a minimum of -7 suggesting that the second site is more extensively cleaved. Importantly, although the 5’-end ratios suggest that the first site is more strongly cleaved, we previously found that the stability of 5’-ends generated from cleavage events does not necessarily correlate with their degree of cleavage^19^.

To broadly compare our new method (Figure 2C-D) to the prior ligation library method, we identified the peaks with a 5’-end ratio ≥ 1000 in MazF’s unfragmented template switch library (n=239). In the fragmented template switch library and the ligation library, these sites featured a sharp drop in read density upstream (Figure S1B). From these 239 cleavage sites, we identified a sequence motif almost identical to that generated by the ligation library method^19^. Additionally, the minimum cleavage ratio within each highly expressed gene was strongly correlated between the ligation and fragmented template switch libraries (R^2^ = 0.70; Figure 2F). This correlation decreased as we decreased the expression threshold used, indicating that cleavage is more difficult to quantify for lower expressed genes (Figure S1C). Taken all together, these results indicate that template switching libraries are a rapid and powerful alternative to ligation libraries that can achieve both single-nucleotide resolution of cleavage sites and quantification of RNA cleavage at each site.

### Endoribonuclease toxins cleave mRNAs with limited specificity

Using our new method, we sought to determine the cleavage specificity of the 9 *E. coli* endoribonuclease toxins that inhibited growth in our system (Figure 1B). We expressed each toxin in a wild-type MG1655 background for 5 minutes from a low-copy plasmid and then generated template switching libraries. As in ligation libraries (Figure 1C), we saw evidence of cleavage across highly expressed regions such as the *atp* operon following expression of all 9 toxins (Figure 3A); the addition of the 5’-end ratios from the unfragmented template switch library resolved many of these regions to specific sites of cleavage (Figure 3B). To globally identify and characterize cleavage sites, we selected 5’-end ratio peaks in the unfragmented libraries that had (i) ≥ 32-fold more signal in the toxin expression sample compared to the empty vector control and (ii) had a log_2_ cleavage ratio ≤ -1 in the fragmented library immediately upstream of the peak. Finally, we required regions to have a high level of expression (≥ 64 reads crossing a given base) in the empty vector sample to ensure we had enough reads to accurately separate cleavage events from noise in poorly-expressed genes. For each toxin, we used the identified cleavage sites (ranging in number from 867 to 3952) to identify sequence motifs directly at the site of cleavage (Figure 3C, left). Because some toxins are ribosome-dependent, we used the annotated reading frames to identify the subcodon position of cleavage within coding regions (Figure 3C, middle). Finally, we plotted the density of cleavage events across coding regions in a 5’-to-3’ direction (Figure 3C, right).

**Figure 3:**
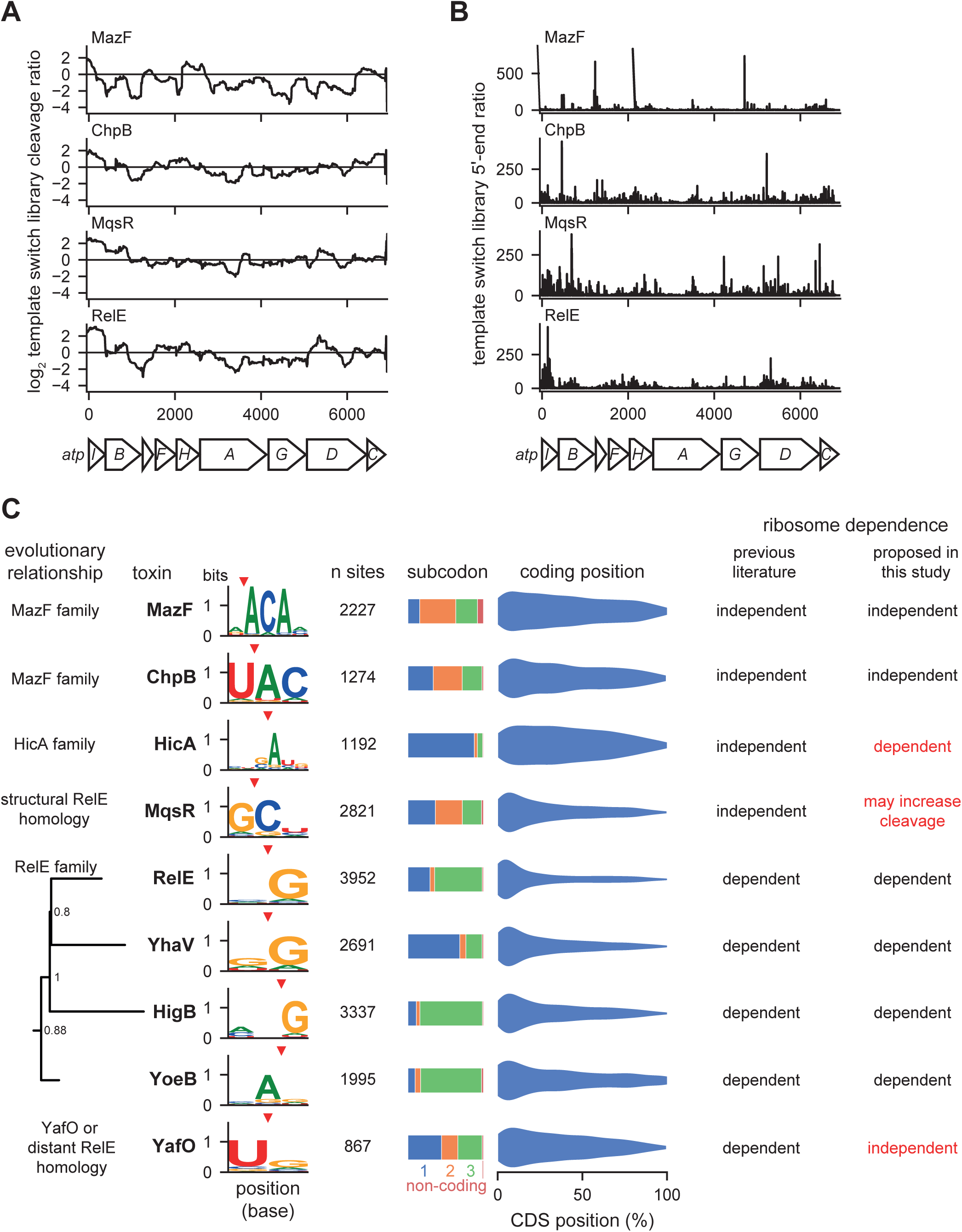
Cleavage specificities of 9 endoribonuclease toxins. **(A-B)** Cleavage profiles (A) and 5’-end ratios (B) from template switch libraries of the highly expressed *atpI-C* region are plotted for the four toxins indicated. Each toxin was expressed for 5 minutes. Data are the mean (A) and geometric mean (B) of 2 replicates for each toxin and empty vector control. **(C)** Summary of cleavage specificities identified for each toxin in expressed non-rRNA regions using a combination of fragmented and unfragmented template switch libraries. Sites (cleavage site + 10 upstream nucleotides) were required to meet a minimum expression threshold of 64 reads at all positions. Sites were called as peaks if the 5’-end ratio was ≥ 32 at the cleavage site with a ≤ 2-fold decrease in read density of the fragmented library in the region upstream of the cleavage site. Nucleotides with ≥ 0.05 bits of information were included in the motifs shown. Evolution tree of RelE family built using 1144 protein sequences with only *E. coli* sequences shown.

Our data indicate that the 9 *E. coli* endoribonuclease toxins examined here have relatively weak cleavage specificities (Figure 3C), consistent with a broad disruption of the transcriptome (Figure 1D). The motifs identified for MazF, ChpB, MqsR, and RelE partially or completely matched previously reported motifs (Table 1). Consistent with the classification of MazF and ChpB as ribosome-independent nucleases, we found that cleavage was not highly dependent on codon position. Additionally, cleavage by MazF and ChpB was seen throughout the coding region of mRNAs, though with some bias to the 5’-end, possibly reflecting differences in RNA secondary structure across coding regions or the fact that cleavage toward the 5’-end of a transcript may destabilize the downstream fragment, reducing the number of subsequent cleavage events detected^26^. MazF and ChpB are both members of the MazF family of nucleases and are more closely related to each other than the other endoribonucleases in *E. coli*. Consistent with this relationship and similarity, MazF and ChpB had similar sequence motifs, with MazF cleaving 5’ of an ACA motif and ChpB cleaving immediately before the AC of a UAC motif.

In contrast to MazF and ChpB, and consistent with its classification as a ribosome-dependent nuclease and in agreement with prior literature^16^, RelE cleavage showed a high dependence on codon position, with most cleavages immediately before a guanosine in the third position of a codon, with a very strong bias in cleavage toward the 5’-end of mRNAs. The toxins YoeB, YhaV, and HigB, which are each members of the RelE family and reported to also be ribosome-dependent, also had (i) a skewed cleavage preference within codons and (ii) a greater preference for the 5’-end of mRNAs. RelE, YhaV, and HigB form a clade and each favors a G just downstream of the cleavage site whereas YoeB has specificity driven by an A upstream of the cleavage site. Though MqsR and YafO have sometimes been included as RelE family members, their cleavage specificity and limited codon position preference diverged from the core RelE family members. Three toxins contradicted the general trends noted for the ribosome-independent and ribosome-dependent toxins. HicA was previously proposed to be ribosome-independent^27^. Although we saw cleavage throughout mRNAs, like with MazF and ChpB, HicA had a strong bias for cleaving before the first position in codons, similar to the ribosome-dependent toxins. MqsR has also been proposed to be a ribosome-independent toxin and has been shown to cleave rRNA precursors^17, 20^. Though we saw no preference with respect to codon position, supporting its ribosome independence, it had a strong bias toward cleaving the 5’-ends of mRNAs, like ribosome-dependent toxins, indicating that the ribosome may play a greater role in its specificity than in other ribosome-independent toxins. Finally, YafO, which was suggested to be ribosome-dependent, had no strong preference with respect to codon position and cleaved throughout mRNAs like the ribosome-independent toxins^28^. These results for HicA, MqsR, and YafO should prompt a reconsideration and further investigation of their interaction with ribosomes and translation.

### tRNA is not a direct target of the endoribonuclease toxins in *E. coli*

In other organisms, endoribonuclease toxins, particularly those in the VapC family, have been shown to target tRNA^15^. Although *E. coli* MG1655 encodes no VapC toxins, we wanted to assess whether the 9 toxins of interest here could cleave tRNA. Mature tRNA is difficult to sequence due to a combination of its small size, structure, and nucleotide modifications, so we first verified that a modified template switch library protocol allowing smaller RNA species could capture tRNA sequences with mature 5’- and 3’-ends by mapping tRNA reads to a consensus tRNA alignment (Figure 4A). Indeed, we observed high read counts across the entire tRNA, with 5’- and 3’-ends corresponding to the expected ends of mature tRNA.

**Figure 4:**
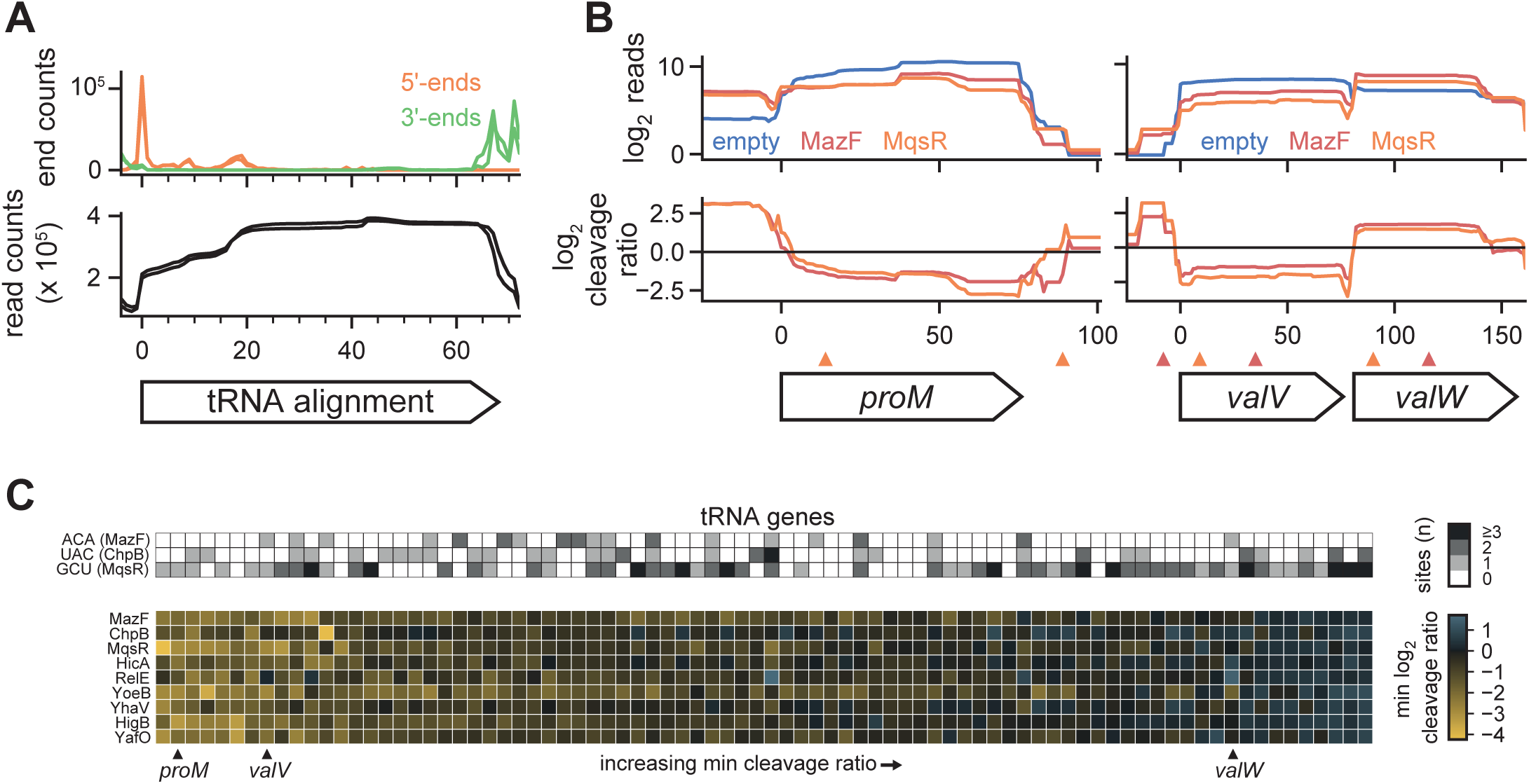
tRNA is not a direct target of *E. coli* endoribonuclease toxins. **(A)** 5’-ends (orange) and 3’-ends (green) of fragments from template switching libraries containing an empty vector (n = 2) were mapped to an alignment of 68 tRNA genes (all tRNA < 78 nt in length except *hisR*; top). The total read counts were also plotted (bottom). **(B)** Read counts (top) and cleavage profiles (bottom) of selected tRNA genes following induction of MazF (red) and MqsR (orange) for 5 minutes compared to the empty vector control. Cleavage sites for MazF (ACA, red) and MqsR (GCU, orange) are plotted as triangles above the genes. **(C)** (Bottom) Heatmap of minimum cleavage ratios in well-expressed tRNAs after 5 minutes of expression of each toxin. (Top) The number of MazF, ChpB, and MqsR cleavage sites within each tRNA is shown as a heatmap. The example tRNAs shown in (B) are marked below.

After expressing each toxin, the read counts for individual tRNAs sometimes changed across the entire gene body, but without producing valleys or cliffs in the cleavage ratio as would be expected for cleavage events (Figure 4B). For example, the toxins MazF and MqsR each altered the expression profiles of the tRNAs encoded by *proM*, *valV*, and *valW*, but without valleys that coincided with the core MazF and MqsR motifs ACA and GCU, respectively. We expanded this analysis to all highly expressed tRNAs (n = 82 of 86), by measuring the minimum cleavage ratio within each tRNA after toxin induction (Figure 4C). Many tRNAs had cleavage ratios close to 0 and in the cases where there were low cleavage ratios, they did not correlate with the presence of cleavage motifs of MazF, ChpB, and MqsR (ribosome-independent toxins with cleavage motifs of at least 3 nucleotides). Further, the changes in tRNA expression observed were broadly correlated across most toxins, supporting a model in which the endoribonucleases indirectly alter the expression, maturation, or recycling of a subset of tRNAs. Taken all together, our results show no evidence of direct tRNA cleavage by *E. coli* toxins but identify common changes in tRNA levels.

### Mature ribosomes are not a target of *E. coli*’s endoribonuclease toxins

Previous work has indicated that MazF family toxins can target rRNAs^18, 29^, although recent studies of *E. coli* MazF and MqsR have shown these toxins primarily target rRNA precursors^19, 20^. To determine if we could identify likely cleavage sites on the rRNA for any other toxins, we generated unfragmented template switch libraries following 30 minutes of toxin expression without ribosome depletion. We then took, using only rRNA regions, the top 30 5’-end ratio peaks and assessed whether the cleavage specificity derived from these peaks matched that found in the wider transcriptome (Figure 5A, S2A). These peaks from rRNAs only matched the characterized cleavage specificity in the cases of MazF, MqsR, and YafO, suggesting that rRNA cleavage by these toxins may be direct. Visual inspection of top cleavage peaks generated by MazF and MqsR showed strong signal unique to each toxin (Figure S2B); YafO peaks were much weaker relative to background suggesting that if cleavage is occurring rRNA is not a common target. Together, these results reaffirm that MazF and MqsR are capable of cleaving rRNA^19, 20^, and suggest that other toxins, with the possible exception of YafO, do not directly cleave rRNA.

**Figure 5:**
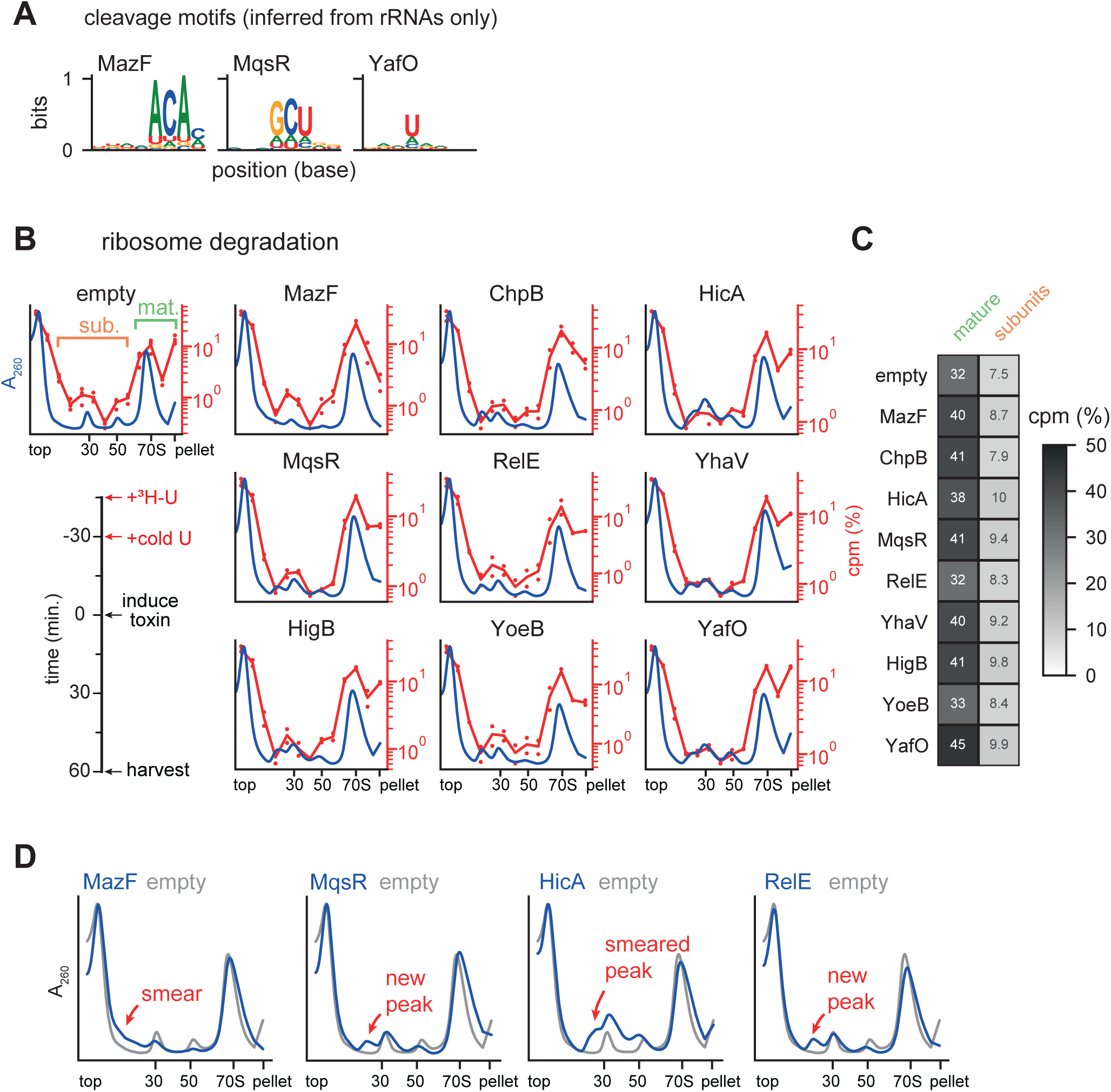
Mature ribosomes are not a target of toxins. **(A)** Cleavage motifs identified from the peaks within rRNA and rRNA precursor regions after 30 minutes of toxin expression. Toxins broadly matching the motif defined in Figure 3C are shown here, with toxins that did not match shown in Figure S2A. **(B)** Sucrose gradients showing effect of toxin expression on mature ribosomes. The timeline of the experiment is shown on bottom left. Average A_260_ values are plotted in blue (empty vector n=8, each toxin n=4). CPMs for ^3^H are plotted on the right axis as a percent of total signal (empty vector n=4, each toxin n=2); dots show the value of each replicate with lines showing the average. **(C)** Summed ^3^H signal from the mature ribosome and precursor fractions from (B). Mature and subunit fractions are defined on the empty vector sample in (B). **(D)** Average A_260_ values for selected toxins compared to the empty vector sample from (B) are enlarged relative to (B) to highlight changes from toxin expression. The location of smears and peaks relative to the empty vector are highlighted.

To better assess whether induction of each endoribonuclease toxin affects mature rRNA, either directly or indirectly, we added ^3^H-uridine to exponential phase cells to label rRNA, chased with unlabeled uridine, induced toxin, and then harvested cells to examine polysomes (Figure 5B, bottom left). This scheme allowed us to assess whether each toxin affected mature, labeled ribosomes. We used sucrose density gradients to separate polysomes (here in the pellet), monosomes, and ribosomal subunits/precursors (Figure 5B). Tritium signal in the mature ribosome fractions was similar in cells expressing toxin or harboring an empty vector, and there was no substantial increase in tritium signal in subunit/precursor fractions for cells expressing toxin (Figure 5C). These results suggest that the toxins do not drive the degradation or disruption of mature ribosomes, or strongly affect the available pool of ribosomes. We conclude that mature ribosome degradation is unlikely to be a major contributor to toxin-dependent growth arrest.

### Endoribonuclease toxins inhibit ribosome biogenesis

After inducing each toxin in the labeling experiment above (Figure 5B), we noticed the appearance of new peaks and smears near the 30S and 50S peaks in the A_260_ traces from our sucrose gradients (Figure 5D). These features were subtle, but highly reproducible and varied between toxins. Because we saw no significant loss in the signal corresponding to mature ribosomes, we inferred that these new peaks arose from disruptions to rRNA synthesis or maturation.

To assess how each toxin impacted the biogenesis of new ribosomes, we induced toxins for 10 minutes and then added ^3^H-uridine without chasing, with cells harvested after an additional 50 minutes of growth (Figure 6A, left). As above, we used sucrose density gradients to separate polysomes (in the pellet), monosomes, and ribosomal subunits (Figure 6A). Relative to an empty vector control, expression of each toxin now substantially increased the incorporation of tritium into the region between the top of the gradient and the monosome peak (Figure 6B). Concomitantly, each toxin also reduced the incorporation of radiolabel into mature ribosomes relative to the empty vector control. Broadly, the fractions containing tritium signal in this experiment were coincident with both the normal subunit peaks observed in the empty vector sample and the aberrant peaks and smears associated with toxin expression (Figures 5D, 6A). Together, these findings indicate that the 9 endoribonuclease toxins examined each disrupt rRNA biogenesis, leading to the accumulation of aberrant rRNA precursors.

**Figure 6:**
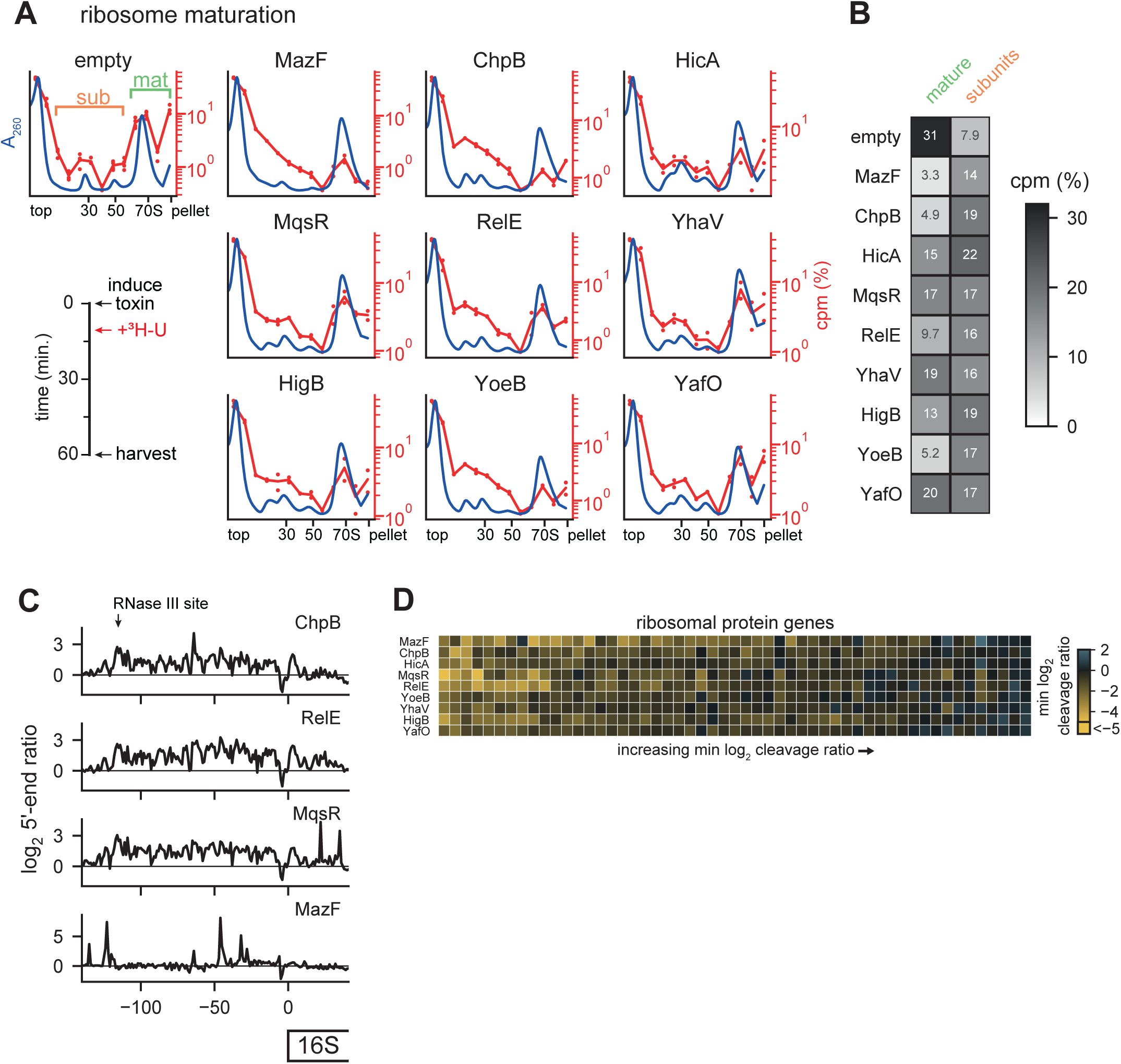
Toxins disrupt ribosome biogenesis. **(A)** Sucrose gradients showing the effect of toxin expression on ribosome biogenesis. Timeline of experiment is shown on bottom left. Average A_260_ values are plotted in blue (empty vector n=8, each toxin n=4). CPMs for ^3^H are plotted on the right axis as a percent of total signal (empty vector n=4, toxin n=2); dots show the value of each replicate with lines showing the average. **(B)** Summed ^3^H signal from the mature ribosome and precursor fractions from (A). Mature and subunit fractions are defined on the empty vector sample in (A). **(C)** Log_2_ 5’-end ratios for four toxins after 30 minutes of expression data was the mean of two replicates each of toxin and empty vector samples. The site of RNase III maturation is labeled in the top plot. **(D)** Heatmap of minimum cleavage ratios for well-expressed (≥ 64 reads at all positions) ribosomal protein genes from fragmented template switch libraries for each toxin after 5 minutes of expression. Data are the mean of two replicates for each toxin and empty vector.

To gather additional evidence, we examined our 5’-end mapping data from the unfragmented template switch libraries, focusing specifically on the 16S and 23S rRNAs after 30 minutes of toxin expression. Here, we found common patterns across all toxins with the exception of MazF. Unlike in mRNAs, where toxins generate clear peaks in the 5’-end ratio (Figure 3B), we observed a broad increase in the 5’-end ratio between the RNase III maturation site and the 5’-end of mature 16S rRNA (Figure 6C); this increase in immature ends supports our conclusion that the new peaks observed on sucrose gradients resulted from improper maturation of rRNA (Figure 6A). The 23S rRNA also showed evidence of aberrant precursors; here there was an increase in 5’-end ratio directly at the RNase III maturation site, again indicating a backlog of unprocessed precursors (Figure S2C). In contrast to the other toxins, MazF’s 5’-end ratio in both regions was dominated by tall peaks like those we observed during cleavage of mRNA (Figure 3B). MazF’s sucrose gradients were also unique compared to the other toxins. While other toxins produced new peaks or peak shoulders near the 30S and 50S, MazF instead decreased the intensity of the two subunit peaks and caused the incorporation of tritium primarily near the top of the sucrose gradient (Figure 5D, 6A). Taken together, these results suggest that MazF may be unique in its strong, direct cleavage of ribosomal precursors and other toxins, including the other ribosome-independent toxins, may inhibit ribosome biogenesis indirectly.

Notably, the same patterns and effects were seen with both ribosome-dependent and ribosome-independent toxins. As the ribosome-dependent toxins like RelE do not directly cleave rRNAs and almost exclusively target translated mRNAs, we conclude that these toxins primarily inhibit ribosome biogenesis by cleaving ribosomal protein transcripts. Thus, our results suggest that these toxins’ inhibition of ribosome biogenesis is likely through inhibition of ribosomal protein synthesis. Since ribosomal proteins are bound while rRNA is synthesized, insufficient ribosomal proteins can lead to defects in folding and proper rRNA maturation^30^. Expression of endoribonuclease toxins have long been associated with decreases in bulk translation, and, using ribosome profiling on cells expressing MazF, we previously linked cleavage sites with ‘traffic jams’ of stuck ribosomes and local translation inhibition^19^. Thus, to determine if ribosomal protein synthesis is inhibited by each toxin’s expression, we measured the cleavage profile minimums in ribosomal protein coding regions (Figure 6D). We observed that toxins inhibit a unique set of ribosomal proteins. The unique set of proteins targeted may lead to different blocks in ribosome maturation and drive the different but reproducible aberrant precursor peaks observed on expression of each toxin. Taken all together, our observations suggest that each toxin creates a set of ribosome precursors incapable of efficiently or properly maturing into normal, 70S particles. Because translation-dependent toxins also produce such precursors, we favor a model in which an improper ratio of ribosomal proteins unique to each toxin’s cleavage specificity leads to a defect in ribosome biogenesis, likely as a primary means of suppressing cell growth.

## Discussion

### Toxins degrade the *E. coli* transcriptome with minimal specificity

A striking feature of toxin-antitoxin systems is that a single strain of bacteria often encodes multiple systems in which the toxins have similar biochemical activities^9^. For endoribonuclease toxins, this observation raises the question of whether individual toxins target the same transcripts to inhibit growth or if individual toxins have specialized functions or specificities. However, to date there has been no systematic, global characterization of all endoribonuclease-based TA systems from a single organism using the same methodology. To do this, we developed a new sequencing pipeline to enable the high-throughput mapping and quantification of RNA cleavage with single nucleotide resolution. The results are in general agreement with our previous study of MazF and other, independent global characterizations of RelE and MqsR^16, 17, 19^. Using our new pipeline, we documented the global patterns of RNA degradation triggered by each of 9 toxins in *E. coli* (Figure 3C).

Each toxin had a short, low complexity cleavage motif, the longest of which was for MazF and ChpB, with high information content at 3 core nucleotides. Notably, differences in nucleotide specificity tracked with evolutionary distance between toxins. Three toxins in our set share clear homology to RelE: YoeB, YhaV and HigB. Of these, YoeB was the most distantly related both evolutionarily and in terms of its cleavage motif, cleaving after an A whereas the others cleaved before a G. MqsR and YafO have been previously suggested to share very distant homology to RelE^31, 32^, and notably both had sequence specificity quite divergent from RelE, YhaV and HigB. ChpB and MazF are homologous and relatively closely related. This similarity was reflected in their similar cleavage specificities, with ChpB’s motif effectively adding a U at position -1 and removing an A at position 3 relative to MazF’s motif. We also noticed that the cleavage specificities of some toxins are similar enough that they target some of the same sites (Figure S3). Assessments of sites that could be cleaved by both MazF and ChpB as well as RelE and HigB showed a greater number of shared sites than would be expected by chance. Given this observation and the generally low complexity of toxin’s motifs, we conclude that toxins are unlikely to specialize in targeting particular transcripts.

Our results support earlier assignments of MazF, ChpB, and MqsR as ribosome-independent toxins as none exhibit strong subcodon bias within translated substrates and MazF and MqsR both show evidence of rRNA cleavage. The related toxins RelE, HigB, YhaV, and YoeB each showed both a subcodon bias and a preference for cleaving the 5’-ends of coding regions, supporting the conclusion that each is a ribosome-dependent toxin^16^. MqsR has also been suggested to also be a ribosome-dependent toxin^32^. In support of this conclusion, we found that MqsR’s cleavage sites were biased towards the 5’-ends of coding regions as with RelE. However, the lack of a subcodon bias and the ability of MqsR to cleave rRNA (Figure 3C, 5A) suggest that translation is not necessary for MqsR cleavage. MqsR’s cleavage pattern is suggestive that strict classification as ribosome-dependent or independent may not be appropriate for some toxins as the ribosome’s activity may play a role in cleavage specificity even without being strictly required for catalysis. HicA was previously characterized as the defining member of a family of ribosome-independent toxins^27^, but it exhibited a strong preference for cleaving immediately before the first position of a codon (Figure 3C). The strength of this codon specificity is hard to reconcile with a completely ribosome-independent model of HicA cleavage. Finally, although YafO was suggested previously to require translation for cleavage, we found that it lacked both a clear subcodon bias and a 5’-end preference (Figure 3C)^28, 31^. Combined with weak signal matching YafO’s cleavage motif from the rRNA (Figure 5A), minimally our results show that YafO behaves differently from other ribosome-dependent endoribonucleases. In sum, our results support a reassessment of the role of the ribosome in substrate specificity in RNase toxins, particularly for MqsR, HicA, and YafO.

### Toxins inhibit ribosome biogenesis through cleavage of ribosome protein transcripts

All of the toxins examined here led to significant disruptions of ribosome biogenesis. We infer that this results primarily through the cleavage of ribosomal protein transcripts, not rRNA directly, as ribosome biogenesis was disrupted even when expressing ribosome-dependent toxins that can only cleave mRNA. Ribosomal proteins bind nascent rRNA as it is transcribed to initiate the maturation of new ribosomes. These ribosomal proteins are required in the correct stoichiometry for efficient folding and maturation of ribosomes so cleavage of their transcripts leads to the disruptions in ribosome biogenesis documented here (Figure 6A-C, S2C). Intriguingly, different toxins created different ‘fingerprints’ of aberrant ribosome precursors (Figure 5D). This observation is consistent with the model that blocking the translation of different sets of ribosomal proteins results in different disruptions of ribosome biogenesis. Future analysis of the RNA and protein content of these precursors might shed light on the stages of ribosome biogenesis that are inhibited and whether they are normal intermediates in ribosome maturation or not. It will also be of interest to determine whether cells in which the relevant antitoxin is restored, and thus that resume growing, are able to redirect these toxin-induced ribosome precursors into mature 70S particles or whether they must first be disassembled or degraded.

### A flexible pipeline for high-throughput bacterial RNA-seq

In this study we developed a new RNA-seq pipeline for identifying the cleavage sites of endoribonuclease toxins (Figure 2A). Combined with improvements to our previously reported rRNA depletion technique^33^, the two un-pooled steps in this pipeline are both 96-well compatible and all subsequent purifications are pooled and rely on magnetic beads to save time and material costs. Because our pipeline uses the first RT step and template switching to integrate PCR amplification handles, it removes many enzymes and reaction steps present in traditional protocols and instead uses only reverse transcriptase and a PCR amplification mix. Together, these measures vastly reduced the cost of sequencing library preparation: a set of 24 libraries with this method costs ∼$10 per library (including rRNA depletion) in enzymes and specialized consumables.

Though we used our pipeline primarily to map cleavage sites, the core protocol can be adapted to capture different types of gene expression data and various RNA species. The most obvious application is to improve the feasibility of large-scale studies of bacterial gene expression; when used with a fragmentation step, the technique can quantify changes in gene expression, even at a sub-gene resolution (Figure 2E-F). Further, by modifying the size-selection protocol, we were able to capture mature tRNA ends (Figure 4A). By pooling early in the pipeline, we needed to conduct only a single gel purification on a barcoded pool of 24 tRNA libraries, making this technique a powerful but simple way to capture tRNA reads. Additionally, we found that our technique also captured 5’-ends arising from native RNA processing during rRNA maturation (Figure 6C, S2B); by combining our approach with enzymatic pre-processing steps already used in other 5’-end mapping protocols^18^, it may be possible to rapidly distinguish transcription start sites and various RNA processing events by selectively sequencing RNA with 5’-triphosphate, 5’-monophoshate, or 5’-OH. Taken together, our approach provides a flexible tool for large scale bacterial RNA-seq studies.

### Concluding remarks

There are an estimated 25,000 ribosomes per cell in actively growing *E. coli* that are capable of translating enough material to double the cell volume every 20 minutes. By inhibiting ribosome biogenesis without degrading mature ribosomes, the endoribonuclease toxins of toxin-antitoxin systems can drive a rapid cessation of growth, while sparing the massive energy investment already made in mature ribosomes, thus avoiding a major fitness cost associated with toxin activation.

For the 9 toxins described here we identified very short, low information cleavage motifs. This limited specificity enables each toxin to drive the degradation of wide swaths of the transcriptome, including ribosomal protein transcripts, leading to a halt in ribosome biosynthesis. However, without knowing the conditions in which toxin-antitoxin systems are normally activated, it remains unclear if ribosome maturation and growth inhibition are their primary target and function, respectively. In principle, endonuclease toxins could inhibit any cellular process requiring synthesis of a protein whose RNA encodes a cleavage site; given their short motifs, the likelihood of inhibiting a process requiring the synthesis of multiple proteins is very high. In the case of bacteria infected with bacteriophage, inhibiting the synthesis of new phage particles may enable populations of bacteria to slow or stop the spread of infection. Indeed, there is growing evidence showing the endoribonuclease activity of toxin-antitoxin systems can both be activated by and defend against bacteriophage^13, 14^. Future exploration of the activation conditions of these toxins will shed light on their role in bacterial survival and stress response.

## Methods

### Growth conditions

*Escherichia coli* was grown in M9 (10x stock made with 64 g/L Na_2_HPO_4_-7H_2_O, 15 g/L KH_2_PO_4_, 2.5 g/L NaCl, 5.0 g/L NH_4_Cl) medium supplemented with 0.1% casamino acids, 0.4% glycerol, 2 mM MgSO_4_, and 0.1 mM CaCl_2_. During experiments, cells were grown at 37°C in either a rotor drum (overnights), an orbital shaker at 200 RPM (bulk cultures for ribosome maturation and degradation), or a Synergy H1 plate reader (BioTek) in 24-well plates using double orbital rotation at 307 CPM (sequencing libraries and toxicity/rescue). All experiments were started from individual colonies following growth overnight on LB (10 g/L NaCl, 10 g/L tryptone, 5 g/L yeast extract) agar plates; colonies were grown up overnight in M9 medium. Antibiotics were used at the following concentrations (liquid/plates): carbenicillin (50 µg mL^-1^ / 100 µg mL^-1^), chloramphenicol (20 µg mL^-1^ / 30 µg mL^-1^).

### Strain and Plasmid construction

Modified pBAD30 plasmids were used for expression of toxins. A sequence containing a ribosome binding site was added between the EcoRI and SacI sites in the MCS of the pBAD plasmid just prior to the toxin’s start codon. Toxin and antitoxin sequences were amplified from the MG1655 *E. coli* chromosome. Antitoxins were inserted into pKVS45 using Gibson assembly. For some TA systems (HicA, YhaV, MqsR, HigB), expression levels of toxin and/or antitoxin were altered by changing the 5’-UTRs of the encoding plasmids to enable rescuable toxicity. All experiments were conducted in MG1655 harboring the toxin-expressing plasmid or both the toxin- and antitoxin-expressing plasmids. Strains are shown in Table 2.

**Table 2.** Strains.

| **Name** | **Genotype** | **Source** |
| --- | --- | --- |
| ML3554 | MG1655 pBAD30 | This study |
| ML3555 | MG1655 pBAD30-mazF | This study |
| ML3556 | MG1655 pBAD30-chpB | This study |
| ML3557 | MG1655 pBAD30-hicA | This study |
| ML3558 | MG1655 pBAD30-yhaV | This study |
| ML3559 | MG1655 pBAD30-mqsR | This study |
| ML3560 | MG1655 pBAD30-rnlA | This study |
| ML3561 | MG1655 pBAD30-relE | This study |
| ML3562 | MG1655 pBAD30-yoeB | This study |
| ML3563 | MG1655 pBAD30-yafO | This study |
| ML3564 | MG1655 pBAD30-yafQ | This study |
| ML3565 | MG1655 pBAD30-higB | This study |
| ML3566 | MG1655 pBAD30 pKVS45 | This study |
| ML3567 | MG1655 pBAD30-mazF pKVS45-mazE | This study |
| ML3568 | MG1655 pBAD30-chpB pKVS45-chpS | This study |
| ML3569 | MG1655 pBAD30-relE pKVS45-relB | This study |
| ML3570 | MG1655 pBAD30-yoeB pKVS45-yefM | This study |
| ML3571 | MG1655 pBAD30-yafO pKVS45-yafN | This study |
| ML3572 | MG1655 pBAD30-mqsR pKVS45-mqsA | This study |
| ML3573 | MG1655 pBAD30-higB pKVS45-higA | This study |
| ML3574 | MG1655 pBAD30-hicA pKVS45-hicB | This study |
| ML3575 | MG1655 pBAD30-yhaV pKVS45-prlF | This study |

### Induction of toxicity and rescue

To measure toxicity and rescue, individual strains harboring each toxin-antitoxin pair were grown from overnight cultures to mid-exponential phase (OD_600_ ∼ 0.25-0.3) in M9 glycerol followed by back dilution to OD_600_ ∼ 0.1 onto a 24-well plate with wells containing either arabinose (0.2% final concentration) or arabinose and anhydrotetracycline (100 ng/mL final concentration) to induce the toxin alone and or the toxin and antitoxin, respectively. Growth was monitored by plate reader set to read OD_600_ every 5 minutes.

### RNA extractions

Cultures (1 mL) were mixed with stop solution (110 µL; 95% ethanol and 5% phenol) and pelleted by centrifugation for 30 seconds at 13,000 RPM on a tabletop centrifuge. Pellets were flash frozen and stored at -80°C. Cells were lysed by adding Trizol (Invitrogen) pre-heated to 65°C directly to pellets followed by 10 minutes of shaking at 65°C and 2000 RPM on a thermomixer (Eppendorf). RNA was extracted from the Trizol mixture using Direct-zol (Zymo) columns following the manufacturer protocol, eluting in 90 µL. Genomic DNA was removed from purified RNA by adding 2 µL of Turbo DNase (Invitrogen) in a 100 µL final volume using the provided buffer and incubating 30 minutes at 37°C. DNase reactions were cleaned up by bringing to 200 µL volume using water and vortexing with 200 µL acid-phenol:chloroform IAA (Invitrogen). After centrifugation, the top layer was extracted and ethanol precipitated with 20 µL 3M NaOAc, 2 µL GlycoBlue (Invitrogen), and 600 µL ice-cold ethanol. After incubation, samples were spun at max speed for 30 minutes at 4°C, washed twice with ice-cold 70% ethanol, and resuspended in water.

### Preparation of ligation RNA-seq libraries

With the exception of MazF data, which was from an earlier study where RNA was extracted from cells expressing MazF for 5 minutes using a similar bulk culture protocol, ligation libraries were prepared as described below. Cells were grown on plates and in overnight cultures as described above with the addition of 0.4% glucose to prevent leaky expression of the toxin from the arabinose inducible promoter. Cultures were back-diluted to 0.02 OD_600_ in 1.5 mL of M9 media with added glucose and grown to OD 0.2 – 0.35 in 24-well plates. Cultures were pelleted by spinning 5 minutes on a benchtop centrifuge at 4000 G. Pellets were washed once with M9 media lacking glucose, respun, and resuspended in M9 media lacking glucose. On a fresh 24-well plate, individual cultures were back-diluted to 0.1 and returned to the plate reader. After 30 minutes of growth, toxin was induced by adding arabinose to 0.2% and cells were returned to the plate reader. After 10 minutes, 1 mL of the culture was removed and used to extract RNA as described above. Ligation RNA-seq libraries were prepared as described previously^19^.

### Preparation of template switch RNA-Seq libraries

Cell cultures were grown and RNA was prepared as described for ligation RNA-seq libraries above. RNA was extracted from samples either 5 minutes or 30 minutes after arabinose induction for all toxins. To deplete rRNA from the 5 minute samples, we used a previously described homebrew rRNA depletion kit as a base protocol and modified it to work in a small volume 96-well format^33^. We first prepared a 100 μM (total molarity, not per oligo) oligo mix of the 19 gram-negative oligos and 2 *E. coli* 5S rRNA specific oligos. Using the calculator and extended protocol (https://github.com/peterculviner/ribodeplete) to calculate volumes and prepare master mixes, we prepared reactions with a 15 μL total volume, an oligo : RNA ratio of 3 and a bead : oligo ratio of 10. Before prepping depletion reactions, streptavidin magnetic beads (21 μL per reaction; NEB) were washed once in an equal volume of 1X SSC and then resuspended in 7.5 μL per reaction 1X SSC with 1 μL per reaction of Superase-In (Invitrogen). Bead/inhibitor mixture was prepped in bulk and aliquoted into individual wells of a 96-well plate and left at room temperature. Next, we prepared depletion reactions. A single depletion reaction had 250 ng of input RNA in a 7.5 μL final volume with 1X SSC, 1 mM EDTA, 0.1 μL oligo mix. Oligos were annealed to the rRNA by placing the depletion reactions on a thermocycler set at 70°C for 5 minutes followed by a gradual (0.1°C/s) ramp down to 25°C. Using a multichannel pipette, the annealing reactions were added to the 96-well plate and pipetted up and down 20 times to mix. After 5 minutes incubation at room temperature, a plastic cover was placed on the plate and it was incubated for 5 minutes at 50°C on a thermocycler. The plate was then removed from the thermocycler and placed on a magnetic rack. Immediately after the reactions clarified (∼30 seconds) a multichannel pipette was used to save 11 µL of supernatant, being careful to not disturb the pellets. The supernatant containing the rRNA-depleted sample was saved in striptubes at -80°C. We used supernatant directly to construct libraries without further purification.

Next, we constructed libraries (extended step-by-step protocol available at: https://github.com/peterculviner/endoribonucmap). Both rRNA depleted samples and total RNA samples were next inputted into the first reverse transcription reaction. For fragmented libraries, 3.75 µL of RNA sample was added to a strip tube containing 1.5 µL of 10x reaction buffer (10x: 500 mM Tris pH 8, 750 mM KCl, 120 mM MgCl_2_) and 1.5 µL of 25 µM reverse transcription primer (since primers were barcoded, a different primer was assigned to each reaction). Fragmentation was accomplished by incubating this high magnesium mixture at 95°C for 3 minutes on a thermomixer and returning to ice. Fragmentation reactions were changed into 15 µL reverse transcription reactions by adding 0.75 µL 10 mM dNTPs, 0.3 µL 100 mM dCTP, 3 µL 100 mM DTT, 3 µL 5M betaine, 0.375 µL Maxima H Minus Reverse Transcriptase (Thermo), 0.75 µL Superase-In (Invitrogen), and 0.075 µL water. To facilitate the large number of reactions, we added these reagents in the form of a master mix and conducted reactions in a 96-well plate. For unfragmented libraries, we followed the above protocol but did not heat reactions after addition of 10x reaction buffer and reverse transcription primer. Reverse transcription reactions were placed on a thermocycler with the following program: 10 minutes at 25°C, 50 minutes at 50°C, and 5 minutes at 85°C. Plates were saved at -80°C after reaction was complete.

Next, reactions with unique barcodes (up to 24 reactions) were combined into a single pool for the rest of the protocol. To ensure we achieved an approximately equal number of reads in our final sequencing run, we conducted the rest of the protocol once using an equal input (2 µL) from each reverse transcription reaction. With this equal pool, we conducted a small scale MiSeq run to determine the contribution of each sample to the final pool; from these read counts, we calculated the correct amount of each reverse transcription reaction to add to achieve an equal number of reads from each reaction.

After combining uniquely barcoded reactions into pools, we brought the volume of each pool to 190 µL with water. To this reaction, we added 10 µL of resuspended AMPure XP beads (Beckman). Resuspended beads were prepared by pelleting 50 µL of magnetic beads into a tube, pelleting them on a magnetic rack, and resuspending them in 10 µL 10 mM Tris pH 8. To the 200 µL sample/bead mixture, we added 200 µL of 20% PEG 8000 (w/v) / 2.5 M NaCl. The reactions were allowed to precipitate for 5 minutes at room temperature before placing on a magnetic rack to pellet. After the tubes clarified (∼5 minutes), we carefully removed the supernatant and washed the pellet twice with 400 µL of fresh 80% ethanol. After removing all ethanol by a quick spin on a benchtop centrifuge, the pellets were allowed to dry at room temperature. After drying, we resuspended each pellet in 10 µL of 10 mM Tris pH 8 / 0.1 mM EDTA by pipetting after removal from the magnetic rack. After allowing the tubes to incubate 5 minutes off the rack, the samples were returned to the rack and the supernatant was transferred to a fresh tube, being careful to not pipette up any magnetic beads.

These pools were next subjected to a second round of reverse transcription to add the 3’ amplification region via template switching. To do this, the 10 µL was added to a strip tube with 4 µL of stock 5x Maxima H Minus Reverse Transcriptase reaction buffer, 0.5 µL of 100 µM template switching oligo, 1 µL of 10 mM dNTPs, 0.4 µL of 100 mM dCTP, 1 µL of Superase-In (Invitrogen), 0.5 µL of Maxima H Minus Reverse Transcriptase (Thermo), and 2.6 µL of water for a final reaction volume of 20 µL. Reactions were placed on a thermocycler with the following program: 10 minutes at 25°C, 30 minutes at 42°C, 5 minutes at 85°C. Following this reaction, samples were stored at -20°C or purified using the same AMPure XP bead-based protocol as described after the first reverse transcription reaction with the exception that reactions were instead resuspended in 20 µL of 10 mM Tris pH 8 / 0.1 mM EDTA.

To identify the correct number of cycles to amplify libraries, we conducted 10 µL trial qPCR reactions on each pool. Trial qPCRs were prepared by mixing 5 µL 2x iTaq Universal SYBR Green Supermix (BioRad), 0.2 µL of any of the i7 amplification primers (100 µM stock), 0.2 µL of the i5 amplification primer (100 µM stock), 3 µL of pre-amplification library, and 2.6 µL of water. Reactions were loaded onto a 384-well plate and run on a Viia 7 qPCR machine using the parameters: 95°C x 5m, (98°C x 30s, 60°C x45s) x 25 cycles. The cycle number where the amplification plot broke ∼½ the maximum signal was used for the final PCR. For the final PCR, we prepared 30 µL reactions using 15 µL 2x KAPA HiFi HotStart ReadyMix PCR Kit (Kapa), 0.6 µL of i7 primer (using a unique i7 primer barcode for each pool, 100 µM stock), 0.6 µL of i5 primer (100 µM stock), 9 µL of pre-amplification library, and 4.8 µL of water. PCR reactions were placed on a thermocycler with the following program: 98°C x 45s, (98°C x 15s, 60°C x 30s, 72°C x 30s) x cycles, 72°C x 60s. After completion, PCR reactions were brought to a 200 µL volume with water and size selected with AMPure XP beads using the following protocol. For each reaction, 50 µL of beads were pelleted on a magnetic rack and resuspended in 100 µL of 20% PEG 8000 (w/v) / 2.5 M NaCl. This mixture was added to each reaction and allowed to incubate 5 minutes at room temperature before returning to the magnetic rack. After the solution clarified, the supernatant (300 µL) was transferred to a fresh tube and the bead pellet (with high MW products) was discarded. Next, 50 µL of beads were pelleted on a magnetic rack and resuspended in 40 µL of 20% PEG 8000 (w/v) / 2.5 M NaCl. This mixture was added to the supernatant to achieve a 340 µL final volume and was allowed to incubate 5 minutes at room temperature before returning to the magnetic rack. After the solution clarified, the supernatant (containing low MW products such as primer dimers) was discarded and the pellet was washed twice with 80% ethanol as described in previous bead purifications. The sample was resuspended in 11 µL of 10 mM Tris pH 8. Libraries were sequenced on either a MiSeq or NovaSeq instrument using a custom read 1 primer. Primers for sequencing library preparation are shown in Table 3.

**Table 3.** Primers. Attached excel spreadsheet.

### Modifications to library protocol for tRNA sequencing

To enable capture of small MW species such as tRNA, we used the protocol described above except for modifying the reaction cleanups between the (1) first and second reverse transcription step and (2) after the second reverse transcription step. For the first clean-up (1), we pooled the tRNA samples as described above but ran the pool on a 10% TBE-Urea gel (Invitrogen) cutting just above a primer dimer band present in a no template control up to ∼150 nt by comparison to a dsDNA ladder. The gel region was crushed and soaked in 400 µL of TE at 70°C for 10 minutes with vortexing. To remove solids from the gel slurry, the mixture was spun through a 0.22 Spin-X filter column (Corning). Small nucleic acids were next precipitated onto AMPure XP beads by resuspending a 100 µL AMPure XP bead pellet in the sample solution and adding 4 parts 1M guanidium HCl in 100% ethanol for each part sample. After incubating 5 minutes at room temperature, samples were loaded on a magnetic rack and allowed to clarify (∼1 minute). After removing the supernatant, the bead pellets were washed twice with 500 µL 80% ethanol and dried and resuspended in 10 µL as described in the standard bead purification protocol above. For the second clean up (2), we modified the AMPure XP bead purification protocol to allow smaller nucleic acids to precipitated. To do this, we resuspended a 50 µL AMPure XP bead pellet in 30 µL of 10 mM Tris pH 8 and added the 20 µL second reverse transcription reaction, 90 µL of 20% PEG 8000 (w/v) / 2.5 M NaCl, and 60 µL of 100% isopropanol. After allowing this to incubate at room temperature for 5 minutes, we placed it on the magnetic rack and proceeded with the rest of the standard bead purification protocol (washes, drying, and resuspension in 20 µL 10 mM Tris pH 8).

### Isotopic labeling of mature and nascent rRNA

Cells containing both toxin and antitoxin vectors were grown in M9 media with a 40 mL final culture volume with a pulse of 5 μCi of [5, 6-^3^H] uridine and 1000-fold excess chase of cold uridine at times indicated. Cells were harvested by centrifugation at 10000 *g* for 1 minute at 4 °C. Cell pellets were washed with once with lysis buffer (20 mM Tris 8, 100 mM NH4Cl, 10 mM MgCl2, 0.5 mM EDTA, and 6 mM β-mercaptoethanol) then respun and resuspended in 300 µL of lysis buffer. To lyse cells, we added 1 µL of Ready-Lyse (Epicenter), 5 µL of Superase-In (Invitrogen), and 2 µL of Turbo DNase (Invitrogen) and incubated the lysis reaction on a thermomixer at 6°C and 500 RPM for 5 minutes. To further promote lysis, we then freeze-thawed cells through 3 cycles of 10 minutes at -80°C and 15 minutes at 6°C and 500 RPM. To remove cellular debris, we then centrifuged lysate for 20 minutes at maximum speed and 4°C on a benchtop centrifuge. The supernatant was loaded onto a 5-20% linear sucrose gradient generated on a Gradient Master (BioComp) instrument in a buffer of 20 mM Tris 8.0, 100 mM NH_4_Cl, and 10 mM MgCl. Samples were centrifuged in an SW41 rotor at 35000 rpm for 4 hours. Gradients were fractionated using the Gradient Master instrument with continuous monitoring of A_260_. 100 μL of each fraction was added to 4 mL of Ecoscint H (National Diagnostics) and ^3^H CPM was measured on a TRI-CARB 4910 TR liquid scintillation counter (PerkinElmer). Measured CPM values were normalized to the volumes of each fraction and to a measurement of CPM quenching across a sucrose gradient standard.

### RelE family phylogeny

To determine the evolutionary relationship between members of the RelE family, a phylogenetic tree of toxin protein sequences was generated. Homologs for each toxin were identified with individual HMMER searches (e value cutoff = 0.01). Resulting sequences were pooled and highly similar sequences were eliminated with CD-HIT (0.7 cutoff, word length = 4)^34^. Resulting 1,144 sequences were aligned using MUSCLE and tree was built using Fasttree^35, 36^. Final tree was pruned to show only E. coli sequences.

### Sequencing data analysis and availability

FASTQ files for each barcode had adapters removed using cutadapt and were mapped to the MG1655 genome (U00096.3) using bowtie2 using the --very-sensitive argument set^37, 38^. FASTQ files for ligation libraries and template switching libraries are available on GEO. For template switch libraries, inline barcodes were also clipped from read 2 and sorted into barcode-specific FASTQ files using cutadapt. Read densities were calculated by identifying the 5’- and 3’-ends of each paired end fragment and adding one count position for all positions aligning to and between the paired reads. For 5’-end sequencing, a count was added only at the 5’-end of read 1. Counts were sequencing depth normalized by calculating size factors based on the total counts mapped to the genome with or without rRNA regions in rRNA undepleted and depleted samples, respectively. Further analyses are described in text and all were accomplished with custom python code and libraries. Code used to generate figures and conduct analyses is available at https://github.com/peterculviner/endoribonucmap. Sequencing data is available at GSE179607 (template switch libraries), GSE144029 (ligation libraries for all toxins except MazF), and GSE107327 (MazF ligation libraries).

## Acknowledgments

We thank C. Guegler for a critical reading of the manuscript. Research was supported by an NIH/NIAID grant (P01AI143575) to S.M.F and an NIH grant (R01GM082899) to M.T.L., who is also an Investigator of the Howard Hughes Medical Institute. The authors have no conflicts to declare.

## Supplemental Figure Legends

**Figure S1:**
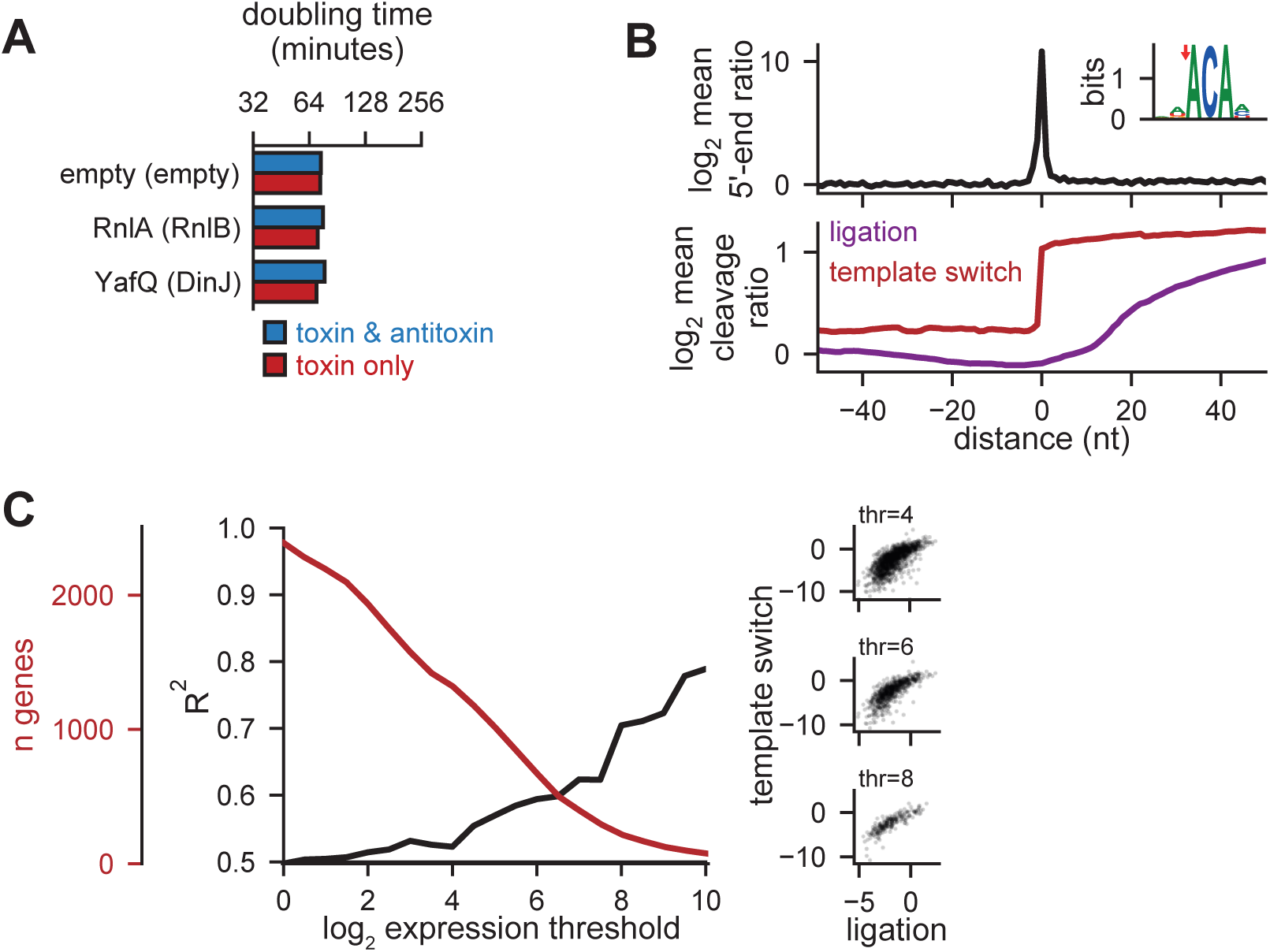
Toxin expression and method development. **(A)** Experiment assessing the toxicity of RnlA and YafQ as in Figure 1B. Doubling times were calculated using time from 90 to 180 minutes after induction. Bars show a single representative sample. **(B)** Mapping the cleavage profiles surrounding the top 239 5’-end ratios following 5 minutes of MazF expression. The average 5’-end ratio at these peaks is shown (top). The motif surrounding these peaks is shown (top, inset). The mean cleavage ratio from the fragmented template switch library and ligation library shown for the region surrounding the top 239 5’-end ratio peaks. **(C)** Comparison of coding region minimum cleavage ratios using template switch libraries and ligation libraries. The R^2^ and number of genes meeting the threshold are plotted across a range of minimum log_2_ expression values (left). Scatter plots of minimum cleavage ratio values using both methods are also shown (right).

**Figure S2:**
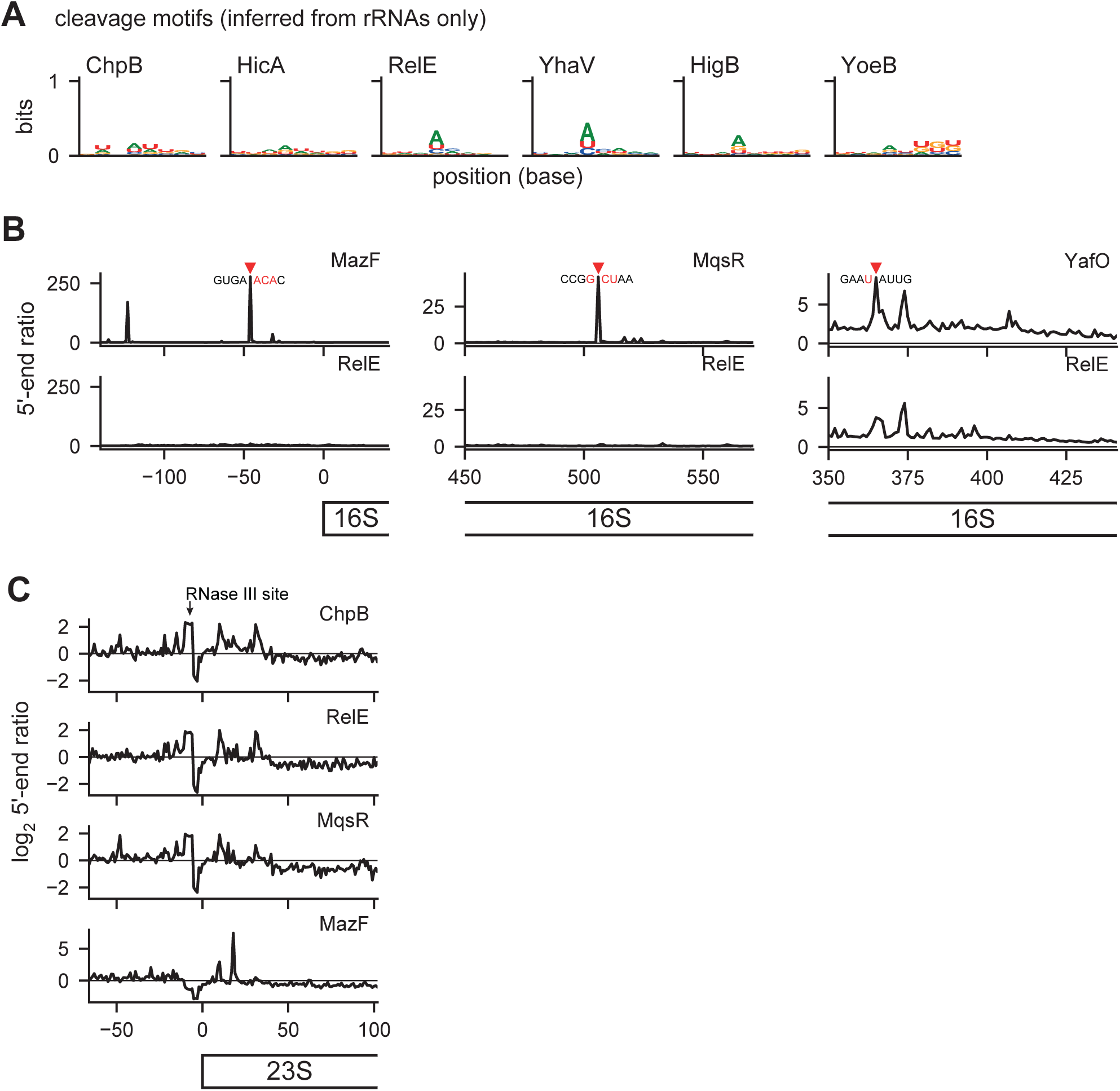
Toxin expression and ribosome cleavage. **(A)** Apparent cleavage motifs identified from rRNA regions for remaining toxins as described in Figure 5A. **(B)** Selected peaks from top 5’-end ratios (geometric mean of n=2) used to define the motifs for MazF, MqsR and YafO. Peaks are shown with red triangles and surrounding 8 nucleotides are shown. Nucleotides matching the core motif are highlighted in red. **(C)** The 5’-end ratios of the 23S loci plotted as in Figure 6C.

**Figure S3:**
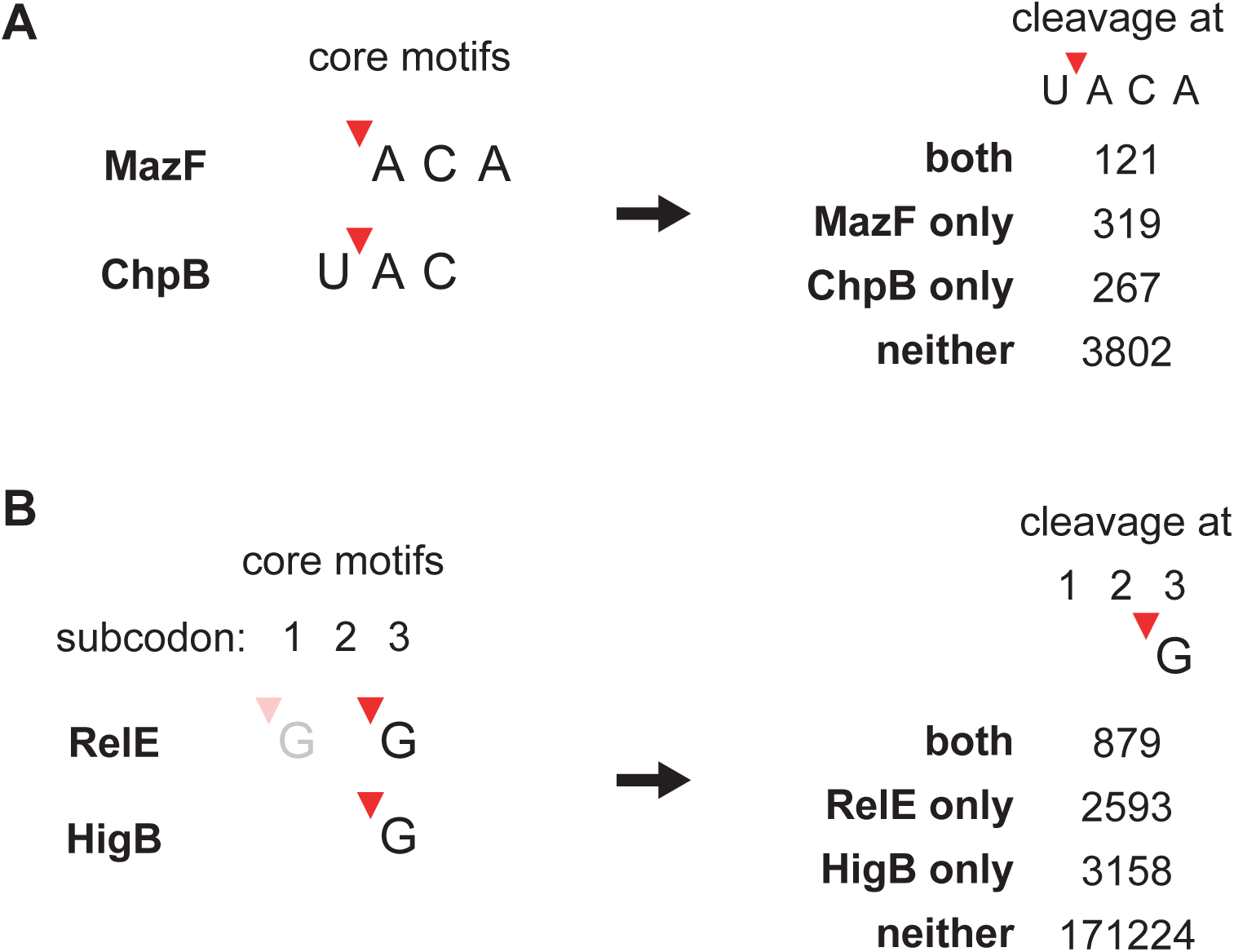
Toxin expression and ribosome cleavage. **(A)** Comparison of core motifs of MazF and ChpB with site of cleavage marked by a red triangle (left). Overlap of cleavage of U^ACA sites by both toxins (right). Cleavage was assessed at sites with ≥ 64 reads in the empty vector sample. Cleavage was defined as a 5’-end ratio ≥ 10 and a cleavage ratio ≤ -1 in the fragmented template switch library. Statistical significance of ‘both’ category was assessed by permuting ‘cleaved’ and ‘uncleaved’ classifications for each toxin across the sites 100,000 times. **(B)** Comparison of RelE and HigB as in Figure S3A. Codon position preferences are shown above and RelE’s lower probability cleavage before subcodon 1 is shown with a greyed-out cleavage motif.

